# Human Nucleolar Protein 7 (NOL7) is required for pre-rRNA transcription and pre-18S rRNA processing

**DOI:** 10.1101/2022.11.08.515626

**Authors:** Mason A. McCool, Carson J. Bryant, Hannah Huang, Lisa M. Ogawa, Katherine I. Farley-Barnes, Samuel B. Sondalle, Laura Abriola, Yulia V. Surovtseva, Susan J. Baserga

## Abstract

The main components of the essential cellular process of eukaryotic ribosome biogenesis are highly conserved from yeast to humans. Among these, the transcription-U3 Associated Proteins (t-UTPs) are a small subunit processome subcomplex that coordinate the first two steps of ribosome biogenesis in transcription and pre-18S processing. While we have identified the human counterparts of most of the yeast Utps, the homologs of yeast Utp9 and Bud21 (Utp16) have remained elusive. In this study, we find NOL7 is the likely ortholog of Bud21. Previously described as a tumor suppressor through regulation of antiangiogenic transcripts, we now show that NOL7 is required for early pre-rRNA stability and pre-18S processing in human cells. These roles lead to decreased protein synthesis, induction of the nucleolar stress response, and defects in cell cycle progression upon NOL7 depletion. Beyond Bud21’s nonessential role in yeast, we establish human NOL7 as an essential UTP that is necessary for both pre-rRNA transcription and processing.

## Methods

### Publicly available expression datasets

Genotype-Tissue Expression (GTEx) unmatched normal and The Cancer Genome Atlas (TCGA) matched normal and tumor expression datasets were obtained through the Xena platform (https://xena.ucsc.edu/) ^1^. RNA-seq by Expectation-Maximization (RSEM) LOG2 fold expression levels for *NOL7* were subtracted from the mean of the overall normal and tumor tissues combined for graphical visualization.

### Sequence alignment and structure analyses

Human NOL7 and *S. cerevisiae* Utp9 and Utp16 protein sequences were used for pairwise alignment using EMBOSS Needle ^2^ to report percent identity and similarity. Vertebrate protein sequences for NOL7 were used for a multiple sequence alignment along with *S. cerevisiae* Bud21 using Clustal Omega ^3^.

*S. cerevisiae* (PDB ID: 5WLC) ^4^ and human (PDB ID: 7MQ8) ^5^ small subunit processome cryo-EM structures were used to analyze Bud21 and NOL7 protein-protein and 5’ETS protein-RNA interactions respectively. Interface residues buried fraction (>0) was considered an interaction using Biojava-structure-6.0.4 for a given position.

### Enrichment analyses

A list of NOL7 interacting proteins was obtained from TheBioGrid ^6^, hu.Map 2.0 ^7^, and a cryo-EM structure of the small subunit processome human pre-A0 cleavage ^5^ (Table S1). This list was used for PANTHER 17.0 gene ontology overrepresentation analysis (Fisher’s Exact test, reported top main categories where fold enrichment > 2, FDR < 0.05, p < 0.05).

### Plasmids

Full-length cDNA for human NOL7 was subcloned into Gateway entry vector pDONR221 (Life Technologies) per manufacturer’s instructions. Site-directed mutagenesis to generate synonymous mutations to render NOL7 resistant to siNOL7 #3 (Table S2) was performed using the Change-IT™ Multiple Mutation Directed Mutagenesis Kit (Affymetrix, 78480) per manufacturer’s instructions. NOL7 siRNA-resistant cDNA was subcloned into the pLX301 vector (Addgene, 25895) containing an N-terminal HA epitope tag with an LR reaction (Life Technologies) per manufacturer’s instructions.

### Cell culture, media, and lentiviral transduction

MCF10A cells (ATCC, CRL-10317) were subcultured in Dulbecco’s modified Eagles’ medium/Nutrient mixture F-12 (Gibco, 1130-032) containing horse serum (Gibco, 16050), 10 μg/mL insulin (Sigma, I1882), 0.5 μg/mL hydrocortisone (Sigma H0135), 100 ng/mL cholera toxin (Sigma, C8052), and 20 ng/mL epidermal growth factor (Peprotech, AF-100-15). HeLa cells (ATCC, CCL-2) cells were grown in DMEM (Gibco, 41965-062) with 10% fetal bovine serum (FBS, Gibco, 10438026). Cell lines were maintained at 37°C, in a humidified atmosphere with 5% CO_2_.

For the high-throughput nucleolar number and 5-EU assays, 3,000 cells/well were reverse transfected in 384-well plates on day 0. For RNA or protein isolation, 100,000 cells/well were seeded into 6-well plates on day 0. For the dual-luciferase assay, 100,000 cells/well were seeded into 12-well plates on day 0.

Lentiviral production and transduction were performed as in ^8^. Lentiviruses were packaged in the HEK293FT cell line by co-transfecting pVSV-G, psPAX2, and modified pLX301 vectors ^9^ containing either a siRNA-resistant version of HA-NOL7 CDS or empty vector in a 1:9:10 ratio (1 μg:9 μg:10 μg for a 10-cm plate) with Lipofectamine 2000 (Life Technologies, 11668019) per manufacturer’s instructions. HeLa cells were transduced with NOL7 or empty vector containing lentivirus under puromycin antibiotic selection (2 μg/mL).

### RNAi

All siRNAs were purchased from Horizon Discovery Biosciences (Table S1). For the 5-EU nucleolar rRNA biogenesis assay of NOL7 ON-TARGETplus pools were used except for the NOL11 positive control where we used the siGENOME SMARTpool siRNAs. For screen validation by deconvolution, the individual ON-TARGET set of four siRNAs that comprised the pool was used. The ON-TARGETplus pools were used in the remaining functional analysis except for northern blotting experiments where siGENOME SMARTpool siRNAs were utilized. Pools of siRNAs were used in all experiments except where indicated that individual siRNAs were tested. siRNA transfection was performed using Lipofectamine RNAiMAX Transfection Reagent (Invitrogen, 13778150) per manufacturer’s instructions with a final siRNA concentration of 20 nM for 384-well plate high-throughput assays and 33 nM for all other assays. For the high-throughput screens, cells were reverse transfected in 384-well plates on day 0 where siRNA controls were added to 16 wells each and siNOL7 was added to 1 well for each replicate. For other assays, cells were transfected 24 h after plating.

### Nucleolar number assay

We followed the ^10^ protocol for counting nucleolar number in MCF10A cells in high-throughput. For siRNA pool treatments, including siONT targeting NOL7, a hit was called based on a stringent cutoff if it produced a mean one-nucleolus percent effect greater than or equal to +3 standard deviations (SD) above the mean percent effect for the entire genome-wide screen (122% effect) (Data S1). For siONT deconvolution, an individual siRNA targeting NOL7 was considered validated if it produced a mean one-nucleolus percent effect greater than or equal to +3 SD above the siNT mean, using the siNT SD within each replicate (Data S2).

### 5-EU incorporation assay

We monitored nucleolar rRNA biogenesis using a high-throughput protocol we developed in the laboratory ^11^. Briefly, following 72 h siRNA depletion, MCF10A cells were treated with 1mM 5-ethynyl uridine (5-EU) for 1 h to label nascent RNA. Then cells were fixed, stained for nucleoli [72B9 anti-fibrillarin antibody ^12^, AlexaFlour 647 goat anti-mouse IGG secondary] and nuclei (Hoëchst), and we performed click chemistry to attach an AF488 azide for 5-EU visualization. Images were acquired with IN Cell 2200 imaging system (GE Healthcare) and a custom CellProfiler pipeline was used for analysis (Data S3).

### Cell Cycle Analysis

We performed cell cycle analysis as in ^13, 14^. Briefly, we analyzed high-throughput screening images from the 5-EU incorporation assay and measured integrated intensity of Hoechst staining per nucleus. The Log2 integrated intensities of each cell were plotted as a histogram where siNT G1 peak was set at 1.0 and siNT G2 peak set at 2.0. siNOL7 treated cell intensities were normalized to siNT. Cell cycle phases were defined as the following normalized Log2 integrated intensities: sub-G1 < 0.75, G1 = 0.75 – 1.25, S = 1.25 – 1.75, G2/M = 1.75 – 2.5, >4n > 2.5.

### Puromycin incorporation Assay

Global protein synthesis was assessed as in ^15^. Following 72 h siRNA depletion, 1 μM puromycin (or 0.5 μM puromycin for Mock 0.5 control) was added to the media for 1 h to label nascent polypeptides. Then we proceeded with western blotting as below.

### Western blotting

Total protein was harvested from cells by scraping followed by PBS rinse. Cells were lysed using AZ lysis buffer (50 mM Tris pH 7.5, 250 mM NaCl, 1% Igepal, 0.1% SDS, 5mM EDTA pH 8.0) with protease inhibitors (cOmplete™ Protease Inhibitor Cocktail, Roche, 11697498001) for 15 minutes at 4°C by vortexing. Lysed cells were spun at 21000 RCF for 15 minutes at 4°C, supernatant was harvested and protein concentration was determined by the Bradford assay (Bio-Rad). Either 50 μg or 25 μg (puromycin blots only) of total protein was separated by SDS-PAGE and transferred to a PVDF membrane (Bio-Rad, 1620177) for blotting. The following primary antibodies were used: α-p53 (Santa Cruz, sc-126), α-puromycin (Kerafast, EQ0001), α-HA (clone 12CA5), and α-β-actin (Sigma Aldrich, A1978), α-POLR1A(Santa Cruz sc-48385), α-FBL (Abcam ab226178), α-UBTF (Santa Cruz sc-13125). For detection of primary antibodies, secondary α-mouse-HRP conjugated antibody (GE Healthcare, Life Sciences NA931) was used. Images were acquired using Bio-Rad Chemidoc (12003153) and analyzed using ImageJ software.

### qRT-PCR analysis

Total cellular RNA was extracted using TRIzol (Life Technologies, 5596018) per manufacturer’s instructions. Prior to cDNA synthesis, all A260/230 values were above 1.7 by NanoDrop (ThermoFisher, ND2000CLAPTOP). cDNA was made from 1μg total RNA using iScript gDNA clear cDNA synthesis kit (Bio-Rad, 1725035) with random primers. qPCR was performed with iTaq Universal SYBR Green Supermix (Bio-Rad, 1725121). The following amplification parameters were used: initial denaturation 95 ºC for 30 s, 40 cycles of 95 ºC for 15 s and 60 ºC for 30 s. Subsequent melt curve analysis was performed to ensure a single product, 95 ºC for 15 s, then gradual (0.3 ºC/15 s) increase from 60 ºC to 94.8 ºC. Gene specific primers were used (Table S3). Amplification of 7SL RNA was used as an internal control and relative RNA levels were determined using comparative CT method (ΔΔCT). Three technical replicates were performed for each biological replicate.

### Dual-luciferase reporter assay

After 48 h of siRNA depletion, MCF10A cells were co-transfected with 1000 ng of pHrD-IRES-Luc plasmid ^16^ and 0.1 ng of a CMV-Renilla constitutive internal control plasmid using Lipofectamine 3000 (Thermo Fisher Scientific, L3000015) per manufacturer’s instructions. Twenty-four h post plasmid transfection (72 h of siRNA depletion), cells were harvested and luminescence was measured by a Dual-luciferase Reporter Assay System (Promega, E1910) per manufacturer’s instructions using a GloMax Discover Microplate Reader (Promega, GM3000). The ratio of pHrD-IRES-Luc / Renilla activity was calculated to control for plasmid transfection efficiency.

### Northern blotting

Total cellular RNA was extracted using TRIzol (Life Technologies, 5596018) per manufacturer’s instructions. For each sample, 3 μg of total RNA was resolved on a denaturing 1% agarose/1.25% formaldehyde gel using Tri/Tri buffer (Mansour and Pestov 2013) and transferred to a Hybond-XL membrane (GE Healthcare, RPN 303S). UV-crosslinked membranes were stained with methylene blue (0.025% w/v) and imaged. Blots were hybridized to ^32^P radiolabeled DNA oligonucleotide probes (5’ETS 5’-CCTCTCCAGCGACAGGTCGCCAGAGGACAGCGTGTCAGC-3’) or (P3 (ITS1) 5’- AAGGGGTCTTTAAACCTCCGCGCCGGAACGCGCTAGGTAC-3’) and detected by phosphorimager (Amersham™ Typhoon™, 29187194). Images were analyzed using ImageJ, ratio-analysis of multiple precursors (RAMP) was performed ^17^.

### BioAnalyzer

For each sample, 100 ng/μL total RNA in nuclease-free water was submitted for Agilent BioAnalyzer analysis, performed by the Yale Center for Genome Analysis.

### Statistical analyses

Statistical analyses were performed using GraphPad Prism 9.3.1 (GraphPad Software). Tests are described in the associated figure legends.

## Introduction

Ribosome biogenesis is an essential and conserved process in all living organisms. In eukaryotes these steps begin in the nucleolus with the transcription of the ribosomal DNA (rDNA) by RNA polymerase I (RNAP1) to synthesize the polycistronic pre-rRNA precursor. This pre-rRNA then undergoes a series of modification, processing, and maturation steps while associating with ribosomal proteins to yield the mature small 40S subunit (18S rRNA) and large 60S subunit (28S, 5.8S, and RNAPIII-transcribed 5S rRNAs) ^18, 19^. Much work has been completed in baker’s yeast, *Saccharomyces cerevisiae* (*S. cerevisiae*), to understand the fundamentals of eukaryotic ribosome biogenesis ^20^. More recent work has highlighted the increased complexities of this process in human cells with new factors, functions, and connections to other cellular processes being uncovered.

The small subunit (SSU) processome is the earliest stable intermediate pre-ribosome complex that forms co-transcriptionally and is marked by the presence of the U3 small nucleolar ribonucleoprotein (U3 snoRNP). Within the SSU processome, there are multiple subcomplexes that are distinguished by their order of assembly ^21-23^. The first subcomplex to form around the pre-rRNA, the transcription U3 associated proteins (t-Utps) or UtpA subcomplex, is required for both transcription and 5’-external transcribed spacer sequence (5’ETS) processing of the primary transcript pre-rRNA. The UtpA subcomplex was first discovered and studied in yeast as a group of essential proteins ^23-25^. Other Utps, including the UtpB and UtpC subcomplexes, associate with the small subunit processome later and thus are only required for pre-18S processing ^26^. Recent work has focused on highlighting the role of the SSU processome’s components through cryo-EM structures of both yeast ^4^ and humans ^5^.

To date, most of the human orthologs of the Utps have been identified except for the t-Utp, Utp9, and a non-essential early associating Utp, Bud21 [called Utp16 in ^24^] (Figure 1A, top). As a t-Utp, Utp9 coordinates both pre-rRNA transcription and pre-rRNA processing ^25^. In part, a direct association with Utp8 is required for its role in pre-18S processing and for an additional role in protein synthesis through direct regulation of mature tRNA nuclear re-export ^27, 28^. Bud21 was first discovered as a non-essential protein required for bud site selection in *S. cerevisiae*. Bud21 is among the set of proteins to associate with the 5’ETS but has not been shown to be a part of any of the SSU subcomplexes in yeast ^21^. Bud21 is also known to be a Ty1 retrotransposon host factor ^29^. Deletion of Bud21 renders increased hypoxia tolerance ^30^ and improved xylose carbon source utilization ^31^, while Bud21 expression is upregulated upon acetic acid challenge ^32^. Overall, not much is known about the molecular functions of Utp9 and Bud21 in ribosome biogenesis, even in the well-studied model organism *S. cerevisiae*.

**Figure 1:**
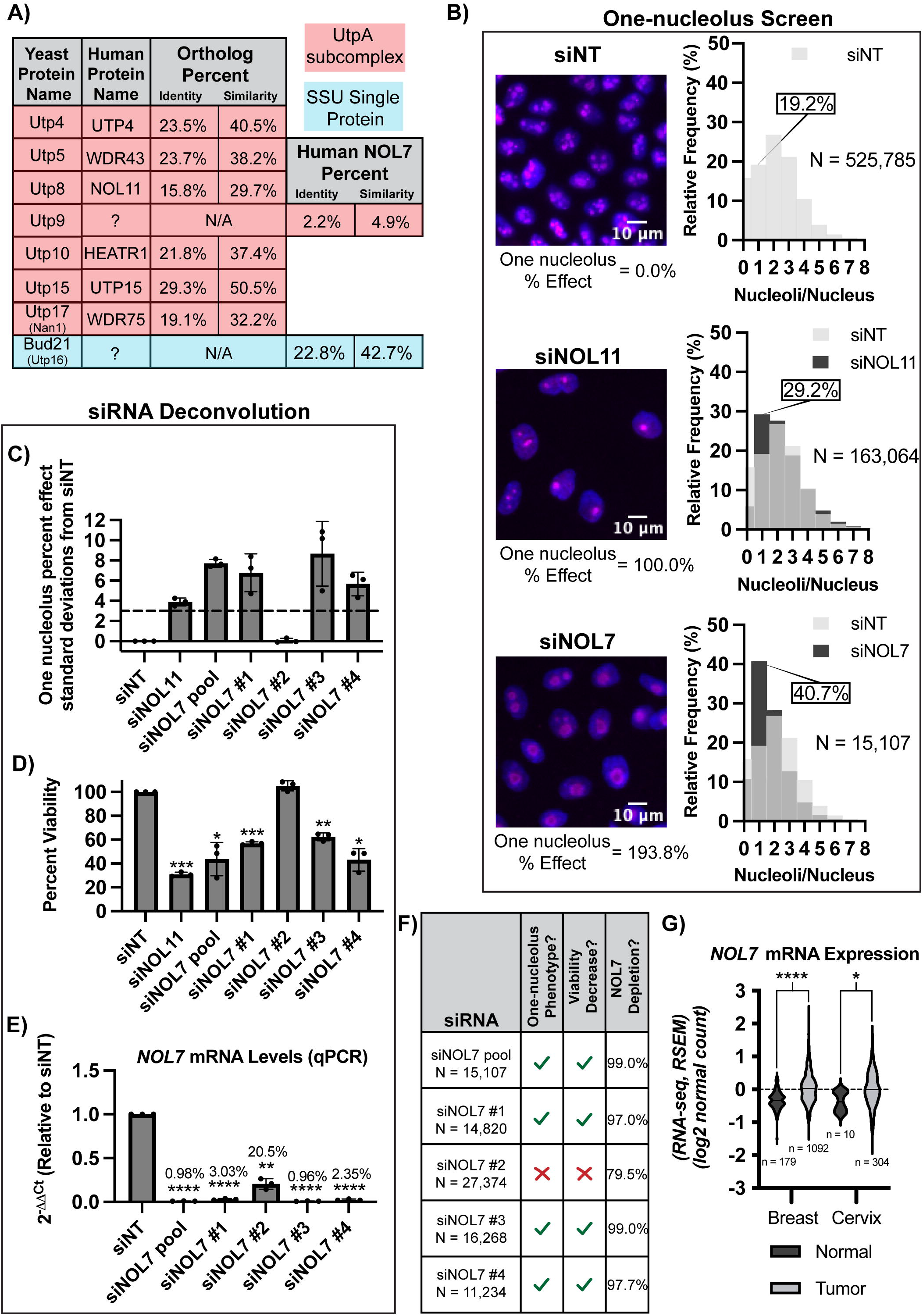
NOL7 is the likely Bud21 (Utp16) ortholog and regulates nucleolar function in human cells. **(A)** Human NOL7 contains the highest sequence similarity to yeast Bud21 (Utp16). Table of yeast and human t-UTP proteins and their orthologs including the yeast t-Utp interacting protein Bud21. Percent similarity and identity of NOL7 with two yeast Utp members that do not currently have a known human ortholog is indicated. **(B)** NOL7 depletion by an siRNA pool reduces nucleolar number in MCF10A cells. (Left) Representative merged images of nucleoli stained with fibrillarin (magenta) and nuclei stained with Hoechst (blue). siNT was used as a negative control (2-3 nucleoli per nucleus), siNOL11 was used as a positive control (1 nucleolus per nucleus). (Right) Histograms of relative frequency of nucleoli per nucleus from quantification of images. Light grey indicates siNT negative control, dark grey indicates either siNOL11 positive control or siNOL7 treated cells. Overlap with siNT indicated by an intermediate grey color. **(C-E)** Deconvolution of the 4 individual NOL7 siRNAs that constitute the NOL7 siRNA pool in MCF10A cells. **(C)** Testing of the 4 individual NOL7 siRNAs for the one nucleolus percent effect. siNOL7 pool and 3 out of 4 individual siRNAs targeting NOL7 produce an increase in the one-nucleolus percent effect in MCF10A cells. 3 biological replicates, plotted mean ± SD. One-nucleolus percent effect is set relative to standard deviations from the negative control, siNT. siNOL11 is a positive control. A 3 standard deviation cutoff (dashed line) was used to consider an siRNA pool treatment to be a hit and for an individual siRNA treatment to pass the deconvolution criteria of producing the one-nucleolus phenotype. **(D)** Testing of the 4 individual NOL7 siRNAs for cell viability. siNOL7 pool and 3 out of 4 individual siRNAs targeting NOL7 reduce MCF10A cell viability. 3 biological replicates, plotted mean ± SD. Cell viability where siNT, negative control, is set to 100%. siNOL11 is a positive control. Data were analyzed by one-way ANOVA with Dunnett’s multiple comparisons test, ** p ≤ 0.01, *** p ≤ 0.001, ** p ≤ 0.01, * p ≤ 0.05. **(E)** Testing of the 4 individual NOL7 siRNAs for *NOL7* mRNA levels. siNOL7 pool and 3 out of 4 individual siRNAs targeting NOL7 reduce *NOL7* mRNA transcript levels greater than 95% in MCF10A cells. qRT-PCR measuring the primary *NOL7* mRNA transcript levels. 2^-^ΔΔ_Ct_ were measured relative to 7SL internal control and siNT negative control sample. 3 technical replicates of 3 biological replicates, plotted mean ± SD. Data were analyzed by one-way ANOVA with Dunnett’s multiple comparisons test, ** p ≤ 0.01, **** p ≤ 0.0001. **(F)** Summary table of siNOL7 pool deconvolution in (C), (D) and (E). Reduction in *NOL7* mRNA levels greater than 95% leads to reduction in nucleolar number and cell viability. N = number of cells analyzed sum of 3 independent replicates. One-nucleolus percent effect passes 3 standard deviations from negative control siNT cutoff from panel E. Viability decreased considered if significance was reached from panel F. The percentage of *NOL7* mRNA level depletion is calculated from panel G. **(G)** Violin plots showing increased expression at the mRNA level for *NOL7* in breast and cervical cancer. Data are from Genotype-Tissue Expression (GTEx) unmatched normal and The Cancer Genome Atlas (TCGA) matched normal and tumor RNA-seq by Expectation-Maximization (RSEM) LOG2 fold expression levels for *NOL7* subtracted from the mean. *NOL7* expression in normal and tumor breast tissue (left), normal and tumor cervical tissue (right). *NOL7* expression in all normal and tumor tissues. Dashed line (set at 0) indicates mean of entire dataset for both normal and tumor expression, black lines indicate mean of individual normal or tumor expression dataset. Data were analyzed by Student’s t-test, **** p ≤ 0.0001, * p ≤ 0.05.

Here, we describe human NOL7 as a likely yeast Bud21 ortholog and establish its function in human cells. NOL7 is required to maintain a normal number of nucleoli in MCF10A cells, an established predictive indicator of proteins that are involved in ribosome biogenesis ^10, 13^. More precisely, NOL7 is required for pre-rRNA transcription, early pre-rRNA stability, and pre-SSU rRNA processing, but not for rDNA promoter activity in human cells. Its depletion leads to decreased mature 18S levels and reduced global protein synthesis, and subsequent induction of the nucleolar stress response. Our results present a new role for NOL7 in human ribosome biogenesis that extends beyond Bud21’s defined role in yeast to emphasize the increasing complexities of early ribosome biogenesis in human cells.

## Results

### NOL7 is the likely yeast Bud21 ortholog

We hypothesized that NOL7 might be the human ortholog to either Utp9 or Bud21 (Utp16) due to its nucleolar localization ^33^ and because of its interactions with components of the SSU processome. We obtained a list of NOL7-associated proteins from high-throughput interactome datasets ^6^ and cryoEM structural analysis ^5^ and performed Gene Ontology overrepresentation tests for both biological processes and cellular components using PANTHER (Log_2_ > 1, p < 0.05) ^34^. NOL7-associated factors’ most enriched categories included positive regulation of rRNA processing (GO:2000234) and transcription by RNA polymerase I (GO:0045943) processes, and the t-UTP complex (GO:0034455) and small-subunit processome (GO:0032040) components (Figure S1A, Table S1). NOL7 protein-protein associations are enriched for factors present in the early SSU processome complex, indicative of a t-UTP subcomplex member.

To address whether Utp9 or Bud21 was the more likely NOL7 ortholog, we performed protein sequence alignments. Interestingly, human NOL7 best aligned in both percent identity and similarity with *S. cerevisiae* Utp16 (22.8% identity, 42.7% similarity) compared to Utp9 (2.2% identity, 4.9% similarity) (Figure 1A). The level of conservation between NOL7 and Bud21 is similar to that of other UTPs (Figure 1A), such as human UTP4 with yeast Utp4 ^35^ and NOL11 with the single-celled eukaryote species *Capsaspora owczarzaki* Utp8 ^36^. We completed a multiple sequence alignment using CLUSTAL Omega ^3^ to visualize the conservation of vertebrate NOL7 with Bud21 across various species (Figure S2). We observed conservation throughout evolution of the Bud21 sequence. Unexpectedly, we were unable to identify NOL7 orthologs in either *Drosophila melanogaster* (^37^ DIOPT Version 8.5 2021, https://www.flyrnai.org/cgi-bin/DRSC_orthologs.pl) or in *Caenorhabditis elegans* (^38^ OrthoList2, http://ortholist.shaye-lab.org/). Furthermore, the overrepresented Gene Ontology categories of interacting proteins were strikingly similar between Bud21 and NOL7 (Figure S1A, B, Table S1). Based on conservation of both sequence and interaction partners, we conclude that NOL7 is most likely the human ortholog of yeast Bud21.

### NOL7 regulates nucleolar function in human cells

NOL7’s functional role in human ribosome biogenesis has not been yet firmly established. As a first pass, we took advantage of our laboratory’s previously established siRNA screening methodology to identify novel regulators of nucleolar function ^10, 13^. Briefly, MCF10A cells normally harbor 2-3 nucleoli per cell nucleus. However, upon depletion of ribosome biogenesis factors, this number decreases to 1 or increases to 5 or more nucleoli on average. Since siRNAs against NOL7 were not present in the original genome-wide siRNA library that we had screened and published ^10^, we specifically depleted NOL7 using siRNAs (si-ONTARGET pool, Horizon Discovery) in high throughput, applying our workflow pipeline. We observed a significant increase in MCF10A cells with 1 nucleolus (40.7% one-nucleolus cells, percent effect = 193.8%) after NOL7 depletion compared to the negative control non-targeting siRNA (siNT) (19.2% one-nucleolus cells, percent effect = 0%). Notably, NOL7 depletion produced a greater proportion of one-nucleolus harboring cells than depletion of the positive control, the t-UTP, NOL11 (29.2% one nucleolus cells, percent effect = 100%) (Figure 1B, Data S1).

To validate NOL7 as a likely hit, we deconvoluted the si-ONTARGET pool targeting *NOL7* to test the ability of each individual siRNA to produce one-nucleolus containing cells and to mitigate possible off-target effects. We observed a reduction in nucleolar number using 3 of the 4 individual siRNAs that targeted NOL7, using > 3 standard deviations from siNT negative control as a stringent cutoff (Figure 1C, Data S2). As expected with inhibition of ribosome biogenesis, this one-nucleolus percent effect produced by individual siNOL7 treatments was inversely correlated with cell viability as measured by number of Höechst-stained nuclei remaining after treatment (Figure 1D). Moreover, the same 3 siRNAs that produced the one nucleolus phenotype also led to the greatest reduction (> 95%) in *NOL7* mRNA levels as measured by qRT-PCR (Figure 1E). Because the other siRNA (siNOL7 #2) did not produce the one-nucleolus phenotype or decrease cell viability, it is possible that ∼20% of normal *NOL7* mRNA levels is enough to maintain normal cellular function (Figure 1F). These deconvolution experiments indicate *NOL7* depletion of >95% is responsible for the hypothesized ribosome biogenesis defects based on the reduction in nucleolar number and in cell viability.

To further demonstrate that the results that we have obtained are due to depletion of NOL7, and not due to an off-target effect of siRNAs, we performed rescue experiments. We made a stable HeLa cell line expressing an siRNA-resistant and N-terminally HA-tagged version of NOL7 (resistant to siNOL7 #3 of the siON-TARGET siRNA pool), and observed that NOL7 mRNA and protein levels depleted by an siRNA can be rescued by overexpressing the si-resistant NOL7. Rescue was detected at both the mRNA level by qRT-PCR and at the protein level by western blotting using an HA antibody (Figure S3). HeLa cells were used for these experiments below to highlight the expected universal role of NOL7 in making ribosomes. Additionally, MCF10A is a non-cancerous and a “near-normal” cell line ^39^, thus using HeLa cells allows for a more direct evaluation if NOL7 is necessary to drive ribosome biogenesis within the context of cancer.

Because NOL7 has been described as a tumor suppressor in previous studies ^33, 40-42^, we analyzed *NOL7*’s mRNA expression levels in breast and cervical cancer (based on cell lines utilized in this study) in Genotype-Tissue Expression (GTEx) unmatched normal and The Cancer Genome Atlas (TCGA) matched normal and tumor samples ^1^. At odds with these previous studies that NOL7 is a tumor suppressor, but consistent with its hypothesized role in making ribosomes, *NOL7* mRNA expression is significantly increased in both breast and cervical cancer tissue compared to normal (Figure 1G). This highlights the relevance of both cell lines used in this study, MCF10A (breast epithelial) and HeLa (cervical cancer). Moreover, we expect NOL7’s role in making ribosomes to be conserved to not only these cell lines but in all human cell types. Concordant with these results, a recent study found that NOL7 expression increases in and helps drive melanoma proliferation and metastasis ^43^. These initial results point towards NOL7 having an important functional role in making ribosomes, and therefore in cell proliferation.

### NOL7 is a component of the SSU processome and necessary for early pre-rRNA stability

Recent cryo-EM structures have offered insight into the location and potential function of Bud21 in yeast ^4^ and NOL7 in humans ^5^ within the SSU processome (pre-A0 5’ETS cleavage). We summarized the interactions of Bud21 and NOL7 from these structures, showing that both proteins make contacts with the 5’ portion of the 5’ETS, the t-UTPs, UTP4 and UTP15, and the U3 snoRNP methyltransferase Nop1/fibrillarin (FBL; Figure 2A). We hypothesized that these direct interactions would give NOL7 the ability to function in pre-rRNA transcription. Although Bud21 has not been found to be required for rDNA transcription in yeast, it is possible that it was overlooked, since Bud21 is non-essential in yeast ^24^.

**Figure 2:**
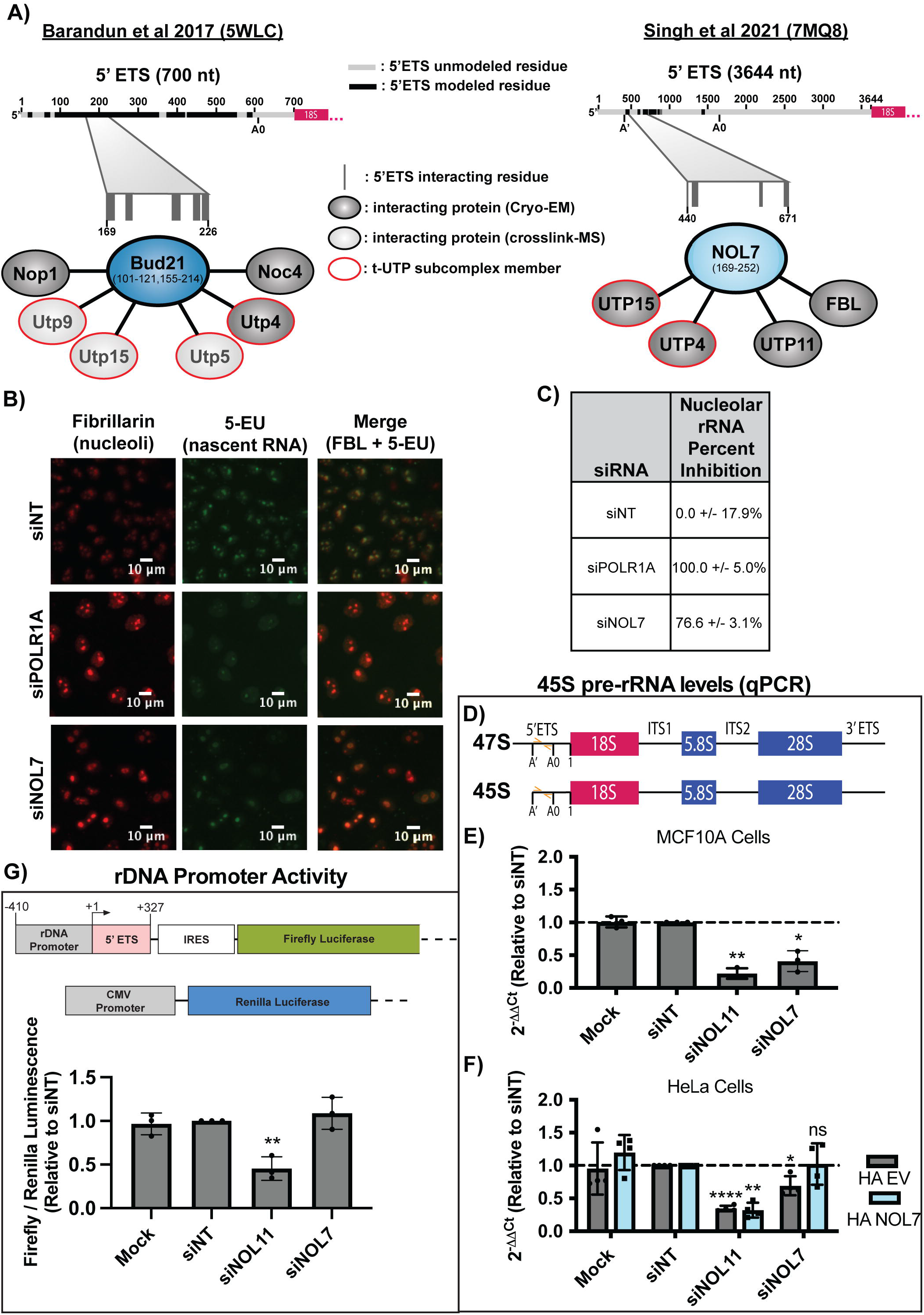
NOL7 is required for rDNA transcription in human tissue culture cells. **(A)** Structure interaction summary of Bud21 (Utp16) and NOL7 within the small subunit processome of (Left) *S. cerevisiae* ^*4*^ (pdb: 5WLC) and (Right) humans ^5^ (pdb: 7MQ8). **(B)** NOL7 siRNA depletion decreases nucleolar rRNA biogenesis. MCF10A cells were depleted with siRNAs for 72 h, then fixed cells were stained for nucleoli (fibrillarin, FBL, red) and click chemistry was performed to conjugate AF488 azide to labeled nascent RNA (5-EU, green). siPOLR1A is a positive control and siNT is a negative control. **(C)** Table of quantification of the percent inhibition of nucleolar rRNA biogenesis after siNOL7 treatment in MCF10A cells. siPOLR1A positive control treatment set at 100%, siNT negative control set at 0%. 3 biological replicates, mean ± SD. **(D-F)** qRT-PCR to measure 45S pre-rRNA levels after NOL7 depletion. **(D)** Schematic of 47S primary and 45S pre-rRNA transcripts with qRT-PCR primer locations indicated (orange). **(E)** NOL7 siRNA depletion reduces 45S pre-rRNA levels in MCF10A cells. qRT-PCR measuring the primary 45S pre-rRNA transcript levels. 2^-^ΔΔ_Ct_ were measured relative to 7SL internal control and siNT negative control sample. siNOL11 is a positive control. 3 technical replicates of 3 biological replicates, plotted mean ± SD. Data were analyzed by one-way ANOVA with Dunnett’s multiple comparisons test, * p ≤ 0.05, ** p ≤ 0.01. **(F)** The reduction of 45S pre-rRNA levels after NOL7 siRNA depletion can be rescued by introduction of an si-resistant version of NOL7. qRT-PCR measuring the primary 45S pre-rRNA transcript in HeLa cells either expressing empty vector (HA EV) or siNOL7 resistant HA-tagged NOL7 (HA NOL7). 2^-^ΔΔ_Ct_ were measured relative to 7SL internal control and siNT negative control sample. siNOL11 is a positive control. 3 technical replicates of 4 biological replicates, plotted mean ± SD. Data were analyzed by one-way ANOVA with Dunnett’s multiple comparisons test, * p ≤ 0.05, ** p ≤ 0.01, *** p ≤ 0.001, ns = not significant. **(G)** NOL7 siRNA depletion does not change rDNA promoter activity. (Top) Schematic of reporter plasmids used: Firefly (pHrD-IRES-Luc, rDNA promoter reporter) and *Renilla* (CMV, transfection control) luciferase plasmids ^16^. (Bottom) Quantification of rDNA promoter activity. Firefly luminescence measured relative to *Renilla* luminescence and siNT negative control. siNOL11 is a positive control. 3 technical replicates of 3 biological replicates, plotted mean ± SD. Data were analyzed by one-way ANOVA with Dunnett’s multiple comparisons test, ** p ≤ 0.01.

To gain insight into NOL7’s potential role in pre-rRNA transcription, we employed an imaging assay developed in our lab to measure nascent nucleolar rRNA biogenesis by 5-ethynyl uridine (5-EU) incorporation and biocompatible click chemistry in MCF10A cells ^11^. By specifically observing the 5-EU signal residing within the nucleolus, we can measure the amount of nucleolar (pre-)rRNA produced over a one-hour time period as a readout of pre-rRNA transcription and stability. Cells depleted of NOL7 had an inhibition of nucleolar rRNA biogenesis of 76.6%, which is almost to the same extent as the positive control, depletion of the largest subunit of RNAP1, POLR1A (100.0% inhibition; Figure 2B, C, Data S3). The level of inhibition observed is consistent with depletion of previously tested factors that are involved in pre-rRNA transcription and processing, including siNOL11, which had an inhibition of nucleolar rRNA biogenesis of 93.8% in our previous study ^11^.

To more precisely define NOL7’s role in pre-rRNA transcription regulation, we measured steady state levels of the primary RNAP1 transcript by qRT-PCR in MCF10A cells depleted of NOL7 (Figure 2D). RNAP1 transcribes the polycistronic 47S pre-rRNA precursor. Cleavage at site A’ in the 5’ETS yields the 45S pre-rRNA precursor ^44^ that is detectable by qRT-PCR (Figure 2D) ^45^. NOL7 depletion resulted in a significant decrease in 45S pre-rRNA levels compared to siNT, similar to the level of depletion of the positive control, NOL11 (Figure 2E). Again, we observed that the decrease in 45S pre-rRNA after NOL7 depletion can be rescued by the expression of an siRNA-resistant version of NOL7, but not an empty vector control in HeLa cells (Figure 2F). Taken together, both the nucleolar rRNA biogenesis assay and the 45S qRT-PCR results point toward NOL7 playing a role in the transcription of the pre-rRNA and its early stability prior to 5’ETS processing, suggesting it is required for optimal RNAP1 transcription in human cells.

Because NOL7 depletion reduces the steady state levels of early pre-rRNA precursors, we tested if this was a result of reduced rDNA promoter activity in siNOL7 treated MCF10A cells. We used a dual-luciferase rDNA promoter (−410 to +327) activity assay ^16^ to assess pre-rRNA transcription after NOL7 depletion in MCF10A cells. In contrast to the results in 2C and 2D, NOL7 depletion did not reduce rDNA promoter activity compared siNT, while the positive control siNOL11 did (Figure 2E). We also measured protein levels of RPA194 and UBTF by western blot to see if NOL7 depletion reduced their abundance and thus pre-rRNA transcription. However, NOL7 depletion did not change either RPA194 or UBTF protein levels (Figure S4).NOL7 is thus required to maintain early pre-rRNA precursor levels, but not through regulation of rDNA promoter (−410 to +327) activity in a reporter system.

### NOL7 is required for U3 snoRNP mediated 5’ETS pre-rRNA processing to produce the small subunit rRNA

t-UTP’s are unique among SSU processome factors as they are required for both the transcription and processing of the pre-18S rRNA ^25, 44, 46^. We tested NOL7’s role in pre-rRNA processing, specifically of the 5’ETS, which is mediated by the U3 snoRNP (Figure 3A) ^47, 48^. Due to NOL7’s interaction with the C/D box snoRNP component, FBL, we hypothesized that depletion of NOL7 could impact the stability of the U3 snoRNA and thus its function in 5’ETS processing. We measured U3 and U8 snoRNA levels by qRT-PCR in MCF10A cells following NOL7 siRNA depletion. U8 is a metazoan-specific C/D box snoRNA involved in large subunit processing ^47, 49^. We observed a significant decrease in U3 snoRNA levels upon NOL7 depletion, but not upon depletion of NOL11. The small changes in U8 snoRNA levels upon NOL7 depletion were not significant. These decreases in U3 snoRNA levels were not associated with any changes in FBL protein levels as measured by western blotting (Figure S4). These results suggest a role for NOL7 in maintaining the stability of the U3 snoRNA.

**Figure 3:**
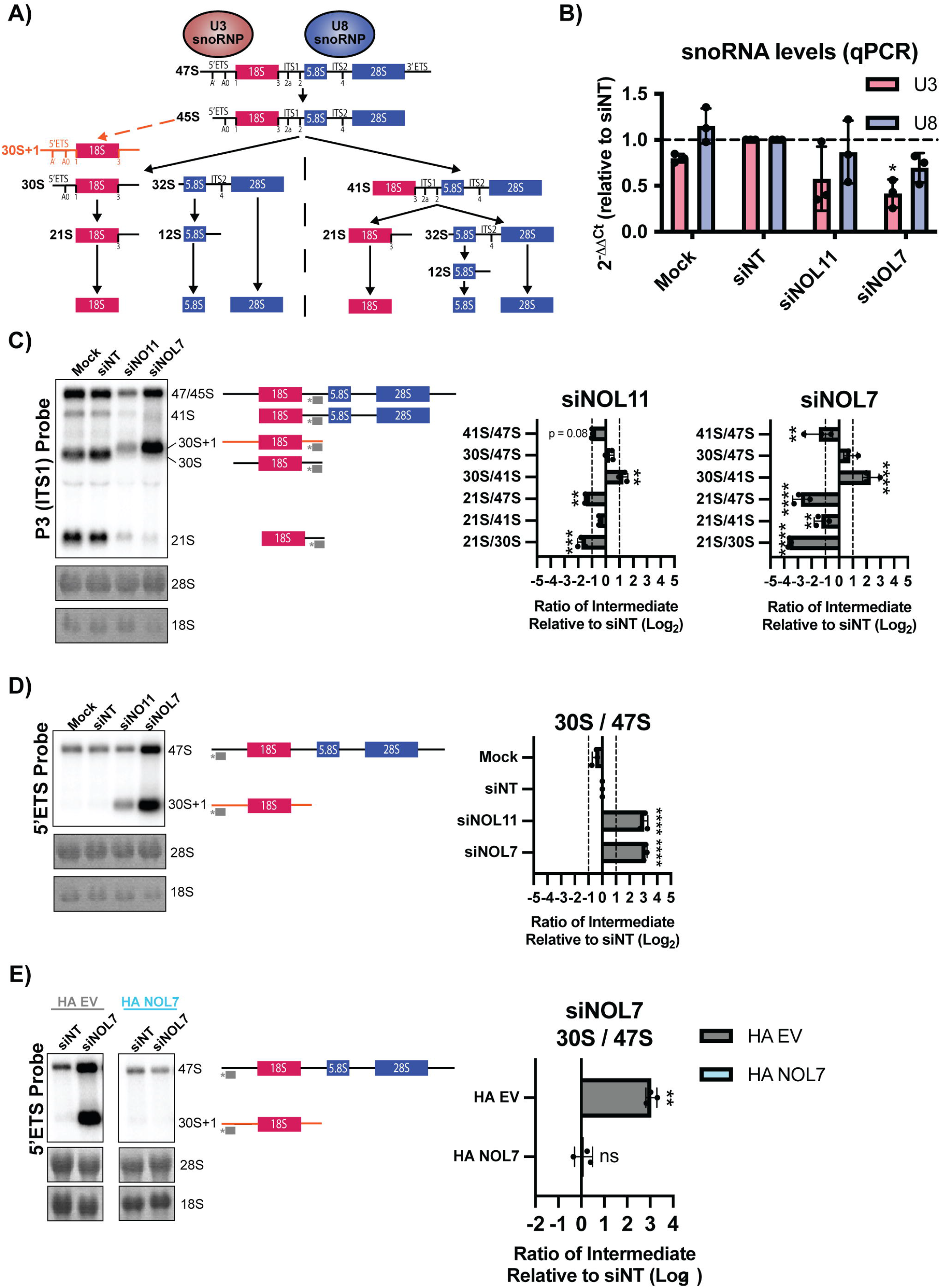
NOL7 is required for U3 snoRNA-mediated small subunit pre-rRNA processing in human tissue culture cells. **(A)** pre-rRNA processing in human cells. The 47S pre-rRNA precursor undergoes a series of modification and processing steps to yield the mature 18S, 5.8S, and 28S rRNAs. Upon inhibition of pre-rRNA cleavage in the 5’ETS, mediated by the U3 snoRNP, accumulation of an aberrant 5’ETS extended 30S+1 precursor occurs (orange). The steps that produce the 18S rRNA (small subunit) are indicated in red, and the steps that produce the 5.8S and 28S rRNAs (large subunit) are indicated in blue. The U8 snoRNA is essential for processing of the rRNAs that make the large ribosomal subunit. **(B)** NOL7 siRNA depletion in MCF10A cells results in reduced U3 snoRNA steady-state levels. qRT-PCR measuring U3 and U8 snoRNA transcript levels in MCF10A cells. 2^-^ΔΔ_Ct_ were measured relative to 7SL internal control and siNT negative control sample. 3 technical replicates of 3 biological replicates, plotted mean ± SD, dotted line at siNT value y = 1. Data were analyzed by one-way ANOVA with Dunnett’s multiple comparisons test, * p ≤ 0.05. **(C)** NOL7 is required for pre-18S rRNA processing in MCF10A cells. (Left) Representative northern blot using a P3 ITS1 probe (indicated in grey) measuring steady state levels of pre-rRNA precursors leading to the 18S rRNA. Methylene blue staining of 28S and 18S was used to show even loading. siNT is a negative control and siNOL11 is a positive control. (Right) Quantification of northern blots using ratio analysis of multiple precursors (RAMP) ^17^ relative to siNT negative control. 3 biological replicates, plotted mean ± SD. Data were analyzed by two-way ANOVA, ** p ≤ 0.01, *** p ≤ 0.001, **** ≤ 0.0001. **(D)** NOL7 is required for 5’ETS cleavage in MCF10A cells. (Left) Representative northern blot using a 5’ETS probe (indicated in grey) measuring steady state levels of 47S and 30S+1 pre-rRNA precursors. Methylene blue staining of 28S and 18S was used to show even loading. siNT is a negative control and siNOL11 is a positive control. (Right) Quantification of northern blots using ratio analysis of multiple precursors (RAMP) ^17^ relative to siNT negative control. 3 biological replicates, plotted Log_2_ (mean ± SD). Data were analyzed by one-way ANOVA with Dunnett’s multiple comparisons test, **** p ≤ 0.0001. **(E)** Accumulation of the 30S+1 pre-rRNA after siNOL7 depletion can be rescued by introduction of an si-resistant version of NOL7 in Hela cells. (Left) Representative northern blot using a 5’ETS probe measuring steady state levels of 47S and 30S+1 pre-rRNA precursors upon NOL7 depletion and si-resistant rescue with either empty vector (HA EV) or HA-tagged NOL7 (HA NOL7) in HeLa cells. Methylene blue staining of 28S and 18S show even loading. siNT is a negative control. (Right) Quantification of northern blots using ratio analysis of multiple precursors (RAMP) ^17^ relative to siNT negative control. 3 biological replicates, plotted Log_2_ (mean ± SD). Data were analyzed by Student’s t-test, ** p ≤ 0.05, ns = not significant.

NOL7 was previously identified in a screen for human proteins required for pre-rRNA processing, where its depletion led to a pre-18S processing defect ^50^. To further validate those findings, we performed northern blots to assess pre-18S rRNA processing in HeLa cells and quantified these results using ratio analysis of multiple precursors (RAMP) ^17^ to detect defects within the processing pathway. First, we probed ITS1 upstream of cleavage site 2 (probe P3) to measure changes in pre-rRNA intermediates leading to the mature 18S rRNA (Figure 3A). Upon depletion of NOL7 and the positive control NOL11, we observed a buildup of 30S pre-rRNA intermediates and a subsequent decrease in 21S pre-rRNA precursors, indicating an inhibition in 5’ETS processing (Figure 3C).

Interestingly, in siNOL7 and siNOL11 treated lanes in the northern blots (Figure 3C) we noticed an upwards shift in the mobility of the 30S pre-rRNA. It is known that upon depletion of factors required for processing of the 5’ETS, there is a build-up of an aberrant 30S+1 intermediate, with an extended 5’ end compared to the 30S intermediate (Figure 3A) ^46, 51^. To more specifically detect this defect, we used a 5’ETS probe upstream of the A’ cleavage site to measure the levels of the 47S and 30S+1 pre-rRNAs. As expected, we observed a significant increase in the 30S+1 intermediate compared to the upstream 47S primary transcript when both NOL7 and the positive control NOL11 were depleted (Figure 3D). This 5’ETS processing defect could be rescued in HeLa cells expressing an siRNA-resistant version of NOL7, but not by an empty vector control (Figure 3E). We conclude from these results that NOL7 is required for 5’ETS pre-rRNA processing in human tissue culture cells.

Since NOL7 is required for pre-18S processing, we tested whether NOL7 depletion would lead to a downstream reduction in the levels of mature 18S rRNA by Agilent BioAnalyzer analysis. There was a significant increase in the 28S/18S rRNA ratio upon NOL7 depletion in MCF10A cells (Figure 4A). More specifically, this increased ratio was due to only decreases in 18S levels rather than increases in 28S levels, indicating NOL7’s role in the maturation of the 18S rRNA (Figure 4B, C). The increase in 28S/18S ratio that was a result of decreases in 18S levels could also be rescued with the introduction of an si-resistant version on NOL7 but not an empty vector control in HeLa cells (Figure 4D-F). NOL7 plays a role in the 5’ETS pre-rRNA cleavage that leads to the 18S rRNA, possibly mediated through its interaction with the U3 snoRNP to maintain its function and stability.

**Figure 4:**
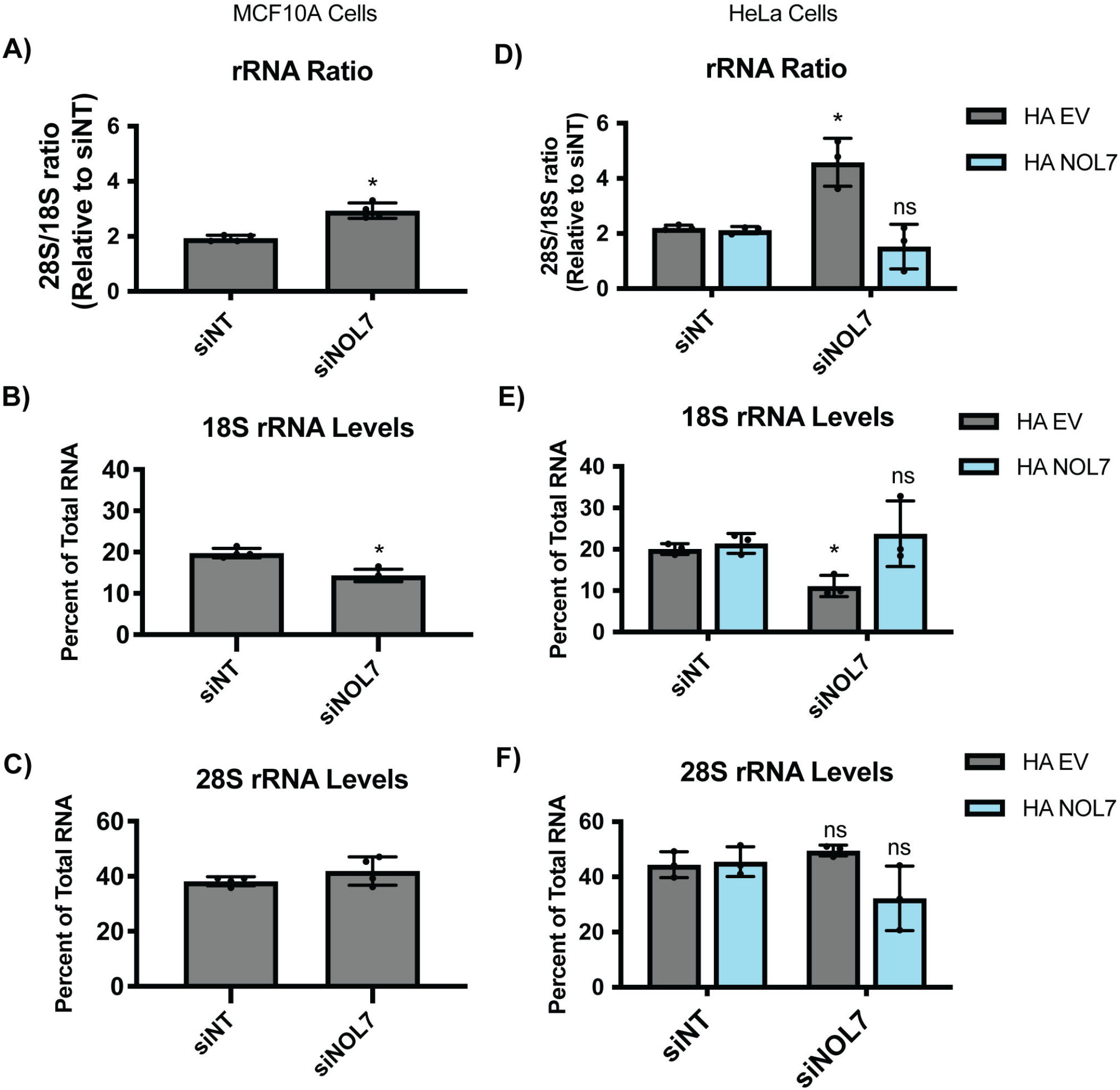
NOL7 is required to maintain mature 18S rRNA levels in human tissue culture cells. **(A)** NOL7 is required for production of the a normal 28S/18S mature rRNA ratio in MCF10A cells. Agilent BioAnalyzer analysis of ratio of mature 28S to 18S rRNAs from MCF10A cells depleted of NOL7. 3 biological replicates, plotted mean ± SD. Data were analyzed by Student’s t-test relative to siNT negative control, * p ≤ 0.05. **(B-C)** NOL7 is small subunit (18S) specific in its role in rRNA maturation in MCF10A cells. **(B)** NOL7 is required for the production of the 18S rRNA. Agilent BioAnalyzer analysis of percent of overall RNA levels for 18S rRNA in MCF10A cells depleted of NOL7. 3 biological replicates, plotted mean ± SD. Data were analyzed by Student’s t-test relative to siNT negative control, * p ≤ 0.05. **(C)** NOL7 is not required for the production of the 28S rRNA. Panel as above (B) except that the large subunit (28S) rRNA was measured. **(D)** siNOL7 increase in 28S/18S mature rRNA ratio can be rescued by introduction of an si-resistant version of NOL7 in Hela cells. Agilent BioAnalyzer analysis of ratio of mature 28S to 18S rRNAs upon NOL7 depletion and si-resistant rescue with either empty vector (HA EV) or HA-tagged NOL7 (HA NOL7) in HeLa cells. 3 biological replicates, plotted mean ± SD. Data were analyzed by Student’s t-test relative to siNT negative control, * p ≤ 0.05, ns = not significant. **(E-F)** siNOL7 small subunit (18S) specific role in rRNA maturation can be rescued by introduction of an si-resistant version of NOL7 in HeLa cells. **(E)** siNOL7 decreases in 18S rRNA levels can be rescued by introduction of an si-resistant version of NOL7. Agilent BioAnalyzer analysis of percent of overall RNA levels upon NOL7 depletion and si-resistant rescue with either empty vector (HA EV) or HA-tagged NOL7 (HA NOL7) in HeLa cells. 3 biological replicates, plotted mean ± SD. Data were analyzed by Student’s t-test relative to siNT negative control, * p ≤ 0.05, ns = not significant. **(F)** siNOL7 and si-resistant version of NOL7 rescue do not change 28S levels. Panel as above in (E) except that large subunit (28S) rRNA was measured.

### NOL7 is required for normal levels of protein synthesis and its depletion leads to induction of the nucleolar stress response

We hypothesized that the role for NOL7 in pre-rRNA transcription, early pre-ribosome stability, and pre-18S processing that we have observed would lead to downstream effects on ribosome function. Therefore, we performed a puromycin incorporation assay to measure changes in global protein synthesis in MCF10A cells depleted of NOL7 ^10, 15^. After siRNA knockdown, cells were treated with 1 μM puromycin, which is incorporated into the nascent polypeptide chain over a time period of 1 h. As expected, depletion of NOL7 led to a significant decrease in global protein synthesis to a similar extent as that of the positive control siNOL11 (Figure 5A, B).

**Figure 5:**
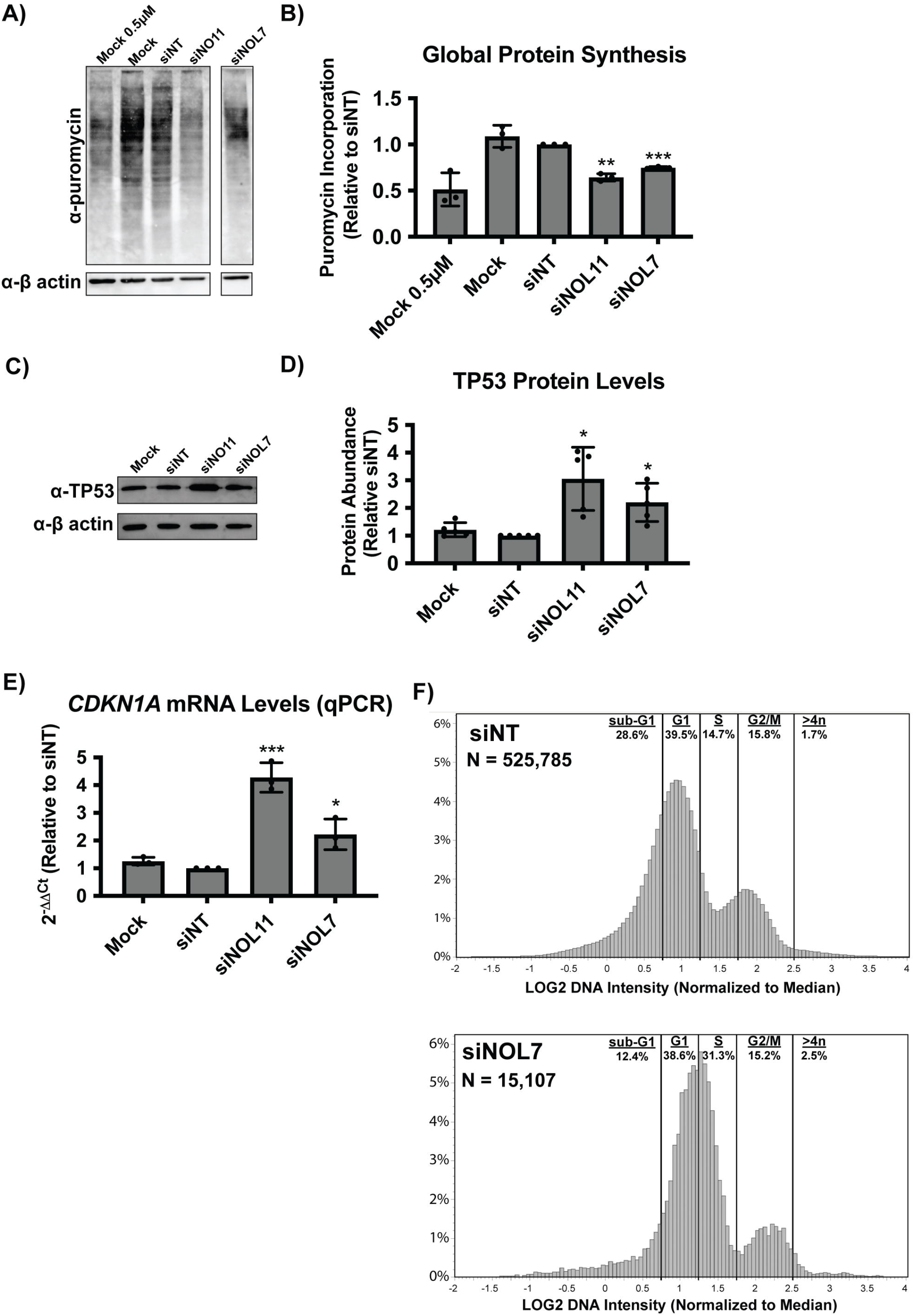
NOL7 siRNA depletion reduces global protein synthesis and causes the nucleolar stress response in MCF10A cells. **(A)** NOL7 siRNA depletion reduces global protein synthesis. MCF10A cells were depleted with siRNAs for 72 h, followed by a 1 h treatment of 1µM puromycin. Total protein was analyzed by western blot. Representative western blot shown where Mock and siNT are negative controls, siNOL11 is a positive control. Mock 0.5µM was included to confirm robustness of the signal quantification. β-actin was used as a loading control. **(B)** Quantification of the decrease in global protein synthesis after NOL7 siRNA depletion in MCF10A cells. Quantification of western blots from (A) normalized to β-actin signal and relative to siNT negative control. 3 biological replicates, plotted mean ± SD. Data were analyzed by one-way ANOVA with Dunnett’s multiple comparisons test, ** p ≤ 0.01. **(C)** NOL7 siRNA depletion leads to increased TP53 protein levels in MCF10A cells. Representative western blot for TP53 shown where Mock and siNT are negative controls, siNOL11 is a positive control. β-actin is a loading control. **(D)** Quantification of the increase in TP53 levels after NOL7 siRNA depletion in MCF10A cells. Quantification of western blots from (C) normalized to β-actin signal and relative to siNT negative control. 5 biological replicates, plotted mean ± SD. Data were analyzed by one-way ANOVA with Dunnett’s multiple comparisons test, * p ≤ 0.05. **(E)** NOL7 siRNA depletion significantly increases *CDKN1A* (p21) mRNA levels in MCF10A cells. qRT-PCR measuring *CDKN1A* mRNA levels. 2^-^ ΔΔ_Ct_ were measured relative to 7SL internal control and siNT negative control sample. siNOL11 is a positive control. 3 technical replicates of 3 biological replicates, plotted mean ± SD. Data were analyzed by one-way ANOVA with Dunnett’s multiple comparisons test, *** p ≤ 0.001, * p ≤ 0.05. **(F)** NOL7 siRNA depletion inhibits cell cycle progression in MCF10A cells. DNA intensity was measured by Hoechst staining. The Log_2_ integrated DNA intensities are plotted as a histogram normalized to siNT where G1 peak = 1.0 and G2 peak = 2.0. Cell cycle phases were the following normalized Log_2_ integrated intensities: sub G-1 < 0.75, G1 = 0.75-1.25, S = 1.25 – 1.75, G2/M = 1.75-2/5, >4n > 2.5. N=the number of cells analyzed.

Because we have defined a role of NOL7 in ribosome biogenesis and in maintaining cell viability, we reasoned that depletion of NOL7 would lead to the nucleolar stress response. As a result of impaired ribosome biogenesis, cells can undergo the nucleolar stress response ^52, 53^. This leads to changes in nucleolar morphology, induction of TP53 levels, downstream induction of *CDKN1A* (p21) expression, and ultimately cell cycle arrest and apoptosis. To check for the induction of the TP53 mediated nucleolar stress response, we measured TP53 levels in MCF10A cells by western blot after NOL7 depletion. As expected, we observed a significant increase of TP53 levels in NOL7 depleted cells (Figure 5C, D). Additionally, since TP53 is a transcription factor for *CDKN1A*, which plays an important role in inducing cell cycle arrest ^54^, we measured *CDKN1A* mRNA levels by qRT-PCR. After depletion of NOL7 or our positive control NOL11, *CDKN1A* mRNA levels were also strikingly increased (Figure 5E). These results are consistent with the observed increase in *CDKN1A* and another cyclin-dependent kinase inhibitor, *CDKN1B* (p27), in melanoma cell lines after NOL7 knockdown ^43^.

We expected that these increases in TP53 and *CDKN1A* levels would result in subsequent cell cycle defects after NOL7 depletion in MCF10A cells. We quantified DNA content of both siNT and siNOL7 treated cells by Hoechst staining from images used in the 5-EU nucleolar rRNA biogenesis assay as described in ^14^. Consistent with previous results in a different human cell line ^43^, we observed an increase in S-phase cells and an accompanying decrease in sub-G1 cells in NOL7 depleted cells compared to siNT (Figure 5F). Taken together, NOL7 is required for maintaining proper ribosome function and cell cycle progression.

## Discussion

We have shown here that the nucleolar protein, NOL7, plays a critical role in early ribosome biogenesis, specifically in pre-rRNA transcription and pre-18S processing, in human cells. siRNA depletion of NOL7 siRNA leads to a reduction in 45S pre-rRNA levels, inhibition of 5’ETS processing, and reduced mature 18S levels. Each of these endpoints were rescued by the introduction of an si-resistant version of NOL7. These defects lead to a downstream reduction in protein synthesis and to the induction of the nucleolar stress response. These results and interactions with other t-UTPs strongly indicate NOL7’s role to be consistent with that of a t-UTP in human ribosome biogenesis.

Some of the roles for NOL7 in human ribosome biogenesis that we have found were expected based on its orthologous yeast protein, Bud21, and some were not. Previously, we had discovered Bud21 as a member of the SSU processome in yeast, but Bud21 was not previously tested for a role in pre-rRNA transcription in this organism ^24, 25^. Bud21 is not essential in yeast and its depletion leads to cold-sensitivity ^24^, a known hallmark of non-essential proteins involved in making ribosomes ^55^. In contrast, NOL7 is essential in most human cancer cell lines as found in the DepMap project (https://depmap.org/) by both shRNA and CRISPR genetic perturbation screens across hundreds of cell lines ^56, 57^. NOL7’s newfound role in ribosome biogenesis is consistent with it being regarded as an essential protein across cancer cell lines. This relevance of NOL7’s function in cancer is further emphasized by cancer cells’ increased reliance on ribosome biogenesis, leading to new therapeutics being developed to target ribosome biogenesis, specifically RNAP1 (i.e. BMH-21) ^58, 59^. It is possible, given the differences in 5’ETS expansion sequences and in interacting proteins, that NOL7 has gained a more impactful function in pre-rRNA transcription and early pre-rRNA stability compared to yeast Bud21 that underlies its essential nature in humans.

While most of our NOL7 findings are consistent with what we know about the function of other t-UTPs, there are some distinct differences that suggest NOL7’s role is not completely aligned with that of all the other t-UTPs. NOL7 is necessary for early pre-rRNA stability and 5’ETS processing that leads to the production of the SSU. More specifically, we discovered that NOL7 is required to maintain 45S pre-rRNA levels and for nucleolar rRNA biogenesis. Each of these assays measures not only pre-rRNA transcription but also pre-rRNA stability. In contrast, in a reporter gene assay for rDNA promoter activity, NOL7 depletion had no effect. Perhaps sequences downstream of the rDNA promoter reporter gene sequences (−410 to +327) are required for NOL7’s function ^16^. Consistent with this, only a fraction of the entire 5’ETS (post A’ cleavage) is modeled in the human SSU processome structure ^5^, none of which is included in our rDNA reporter gene. Additionally, NOL7 is important for maintaining steady-state levels of the U3 snoRNA, with which it interacts through FBL based on the SSU processome structure ^5^.

Although NOL11 depletion did not recapitulate the same results, it remains to be elucidated how other t-UTPs and SSU processome factors alter the stability of early pre-ribosomes and the U3 snoRNA during transcription and the first steps of pre-rRNA processing in human cells.

Our discovery of NOL7’s essential role in making ribosomes is at odds with several previous studies on NOL7’s role as a tumor suppressor. These studies focused on NOL7’s role in post-transcriptional regulation of mRNAs, more precisely mRNA stabilization through direct interactions with mRNAs and their processing machinery within the nucleus ^33, 40-42^. These interactions lead to a tumor suppressor effect due to preferential stabilization of antiangiogenic transcripts, of which TSP-1 has been validated ^42^. It is worth noting that within our PANTHER overrepresentation analysis of Bud21 interacting proteins, but not for the shorter list of known NOL7 interactors, we discovered an enrichment of GO biological processes consistent with this other NOL7 function including: intracellular mRNA localization (GO:0008298), positive regulation of mRNA catabolic process (GO:0061014), regulation of mRNA stability (GO:0043488), and RNA transport (GO:0050658) (Figure S1B). Similarly, there is precedent for other ribosome biogenesis factors having functions in cellular processes outside the nucleolus, including nucleolin and FANCI, in which their roles are regulated by their interacting partners and subcellular localization ^8, 60^. It is unsurprising that the ubiquitous process of making ribosomes has cross-talk mechanisms to coordinate itself with other cellular processes. Our data and ^43^ strongly suggest NOL7’s role to be important for tumor growth and progression, but previous data indicates a role opposing this as well. It remains to be further explored to what extent and in which contexts NOL7 functions outside of ribosome biogenesis, particularly through modulation of mRNA subsets.

We discovered that depletion of NOL7 reduces cell viability and cell cycle progression, most likely as a result of the nucleolar stress response. These results agree with ^43^ in which NOL7 depletion increases *CDKN1A* levels and leads to cell cycle S-phase arrest in melanoma cell lines. While the nucleolar stress response occurs as a result of impaired ribosome biogenesis, it is also plausible that NOL7 could act in a more direct manner to modulate the cell cycle. Regulation of cell cycle (GO:0051726) was one of the top overrepresented biological processes categories among NOL7 interacting proteins (Figure S1A). While our data and analysis emphasize NOL7’s role in ribosome biogenesis, it remains to be studied if NOL7’s interaction with these proteins involved in the cell cycle has a more direct impact on cell cycle regulation and overall cell viability.

NOL7 is another example of the increased complexities of ribosome biogenesis in humans as compared to yeast. We have demonstrated this through NOL7’s role in pre-rRNA transcription, processing, and early pre-rRNA and U3 snoRNA stabilities. It is also exemplified by NOL7’s other known function outside the nucleolus in regulation of nuclear mRNA stability ^33, 40-42^, adding even more layers of complexity. These results and future studies will certainly help uncover the finer details of human ribosome biogenesis, specifically pre-rRNA transcription and SSU processome function, that will lead to better understanding of therapeutic design options in diseases such as cancer and ribosomopathies.

