## Supplementary Figures for "Human Nucleolar Protein 7 (NOL7) is required for pre-rRNA transcription and pre-18S rRNA processing"

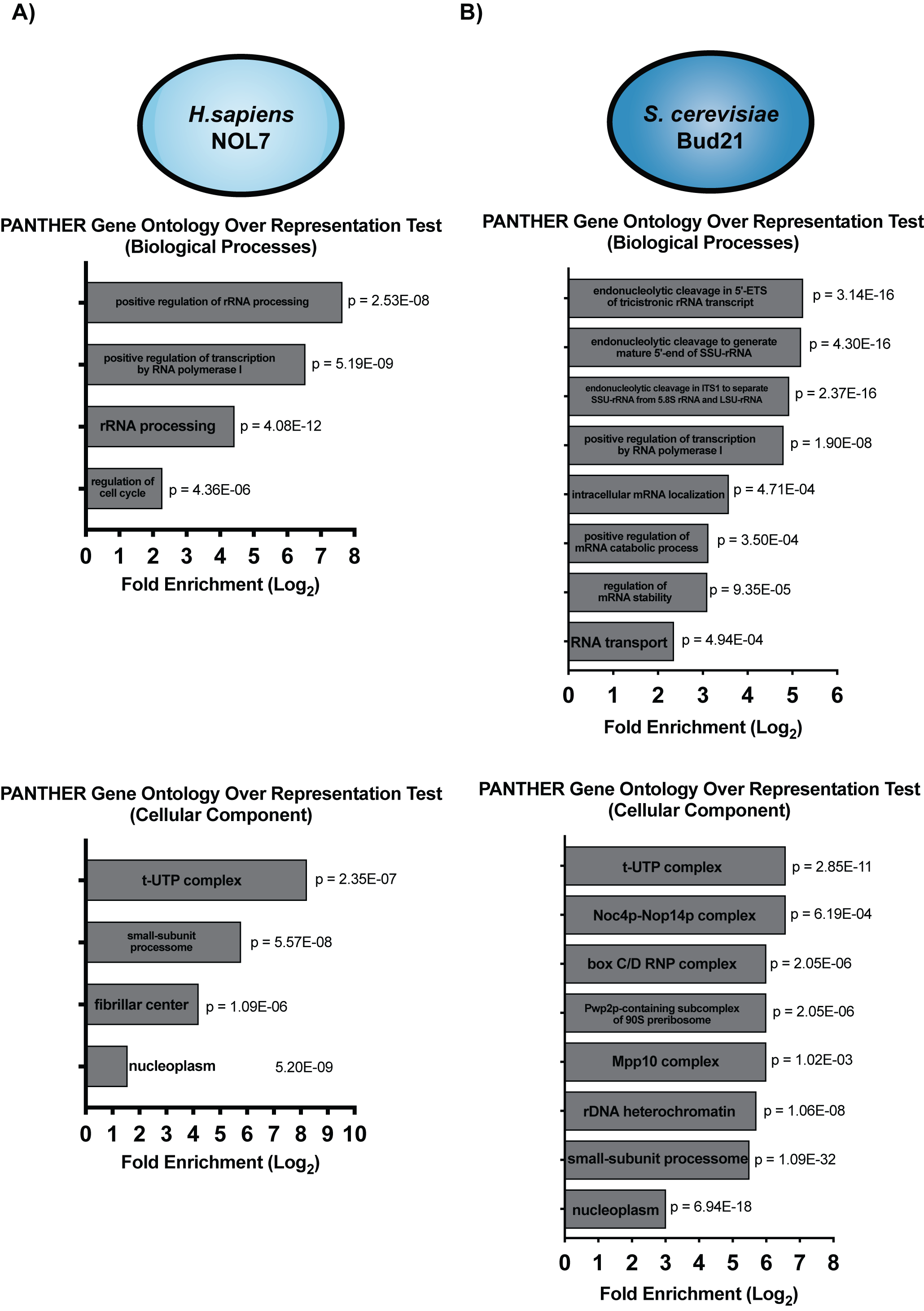

**Figure S1**: Human NOL7 and yeast Bud21 associate with similar proteins. **A)** A list of NOL7 interacting proteins was compiled from TheBioGrid (Oughtred et al. 2021) and direct contacts from human pre-A0 cleavage small subunit processome cryo-EM structure pdb: (Singh et al. 2021). A gene ontology overrepresentation test for Biological Processes (top) and Cellular Component (bottom) was completed using PANTHER 17.0. Data were analyzed by Fisher’s Exact test, fold enrichment > 2, FDR < 0.05, p < 0.05. **B)** Same analysis as in **A)** except for Bud21 interacting proteins from TheBioGrid and direct contacts from *Saccharomyces cerevisiae* small subunit processome cryo-EM structure pdb: 5WLC (Barandun et al. 2017) were used.

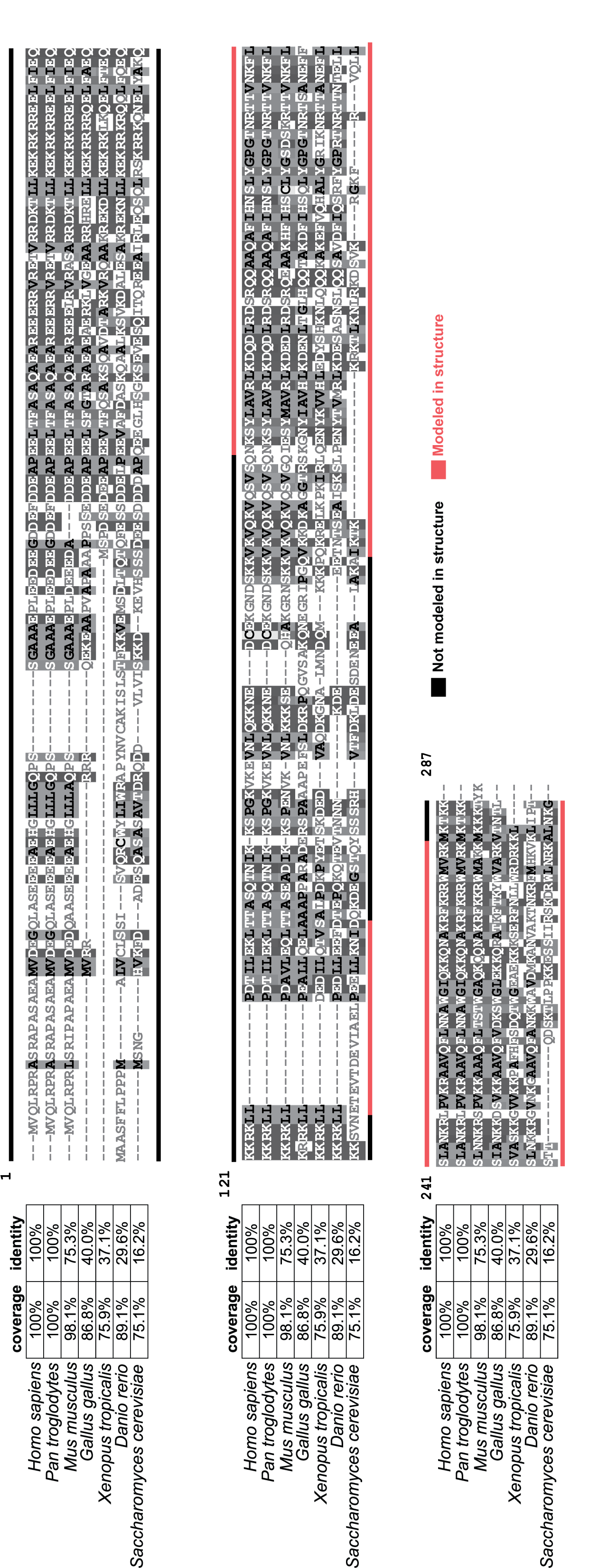

**Figure S2**: The Bud21 protein sequence is conserved from *Saccharomyces cerevisiae* to vertebrate NOL7. Multiple protein sequence alignment of NOL7 in different vertebrate species with *S. cerevisiae* Bud21 using Clustal Omega (Sievers et al. 2011). Percent coverage and identity values shown in table on left. Consensus disorder (not in modeled in cryo-EM structure, black) and U3 snoRNA associated (modeled in cryo-EM structure, red) indicated with a line either below for Bud21 within yeast (Barandun et al. 2017) or above for human (Singh et al. 2021) small subunit processome structures, respectively.

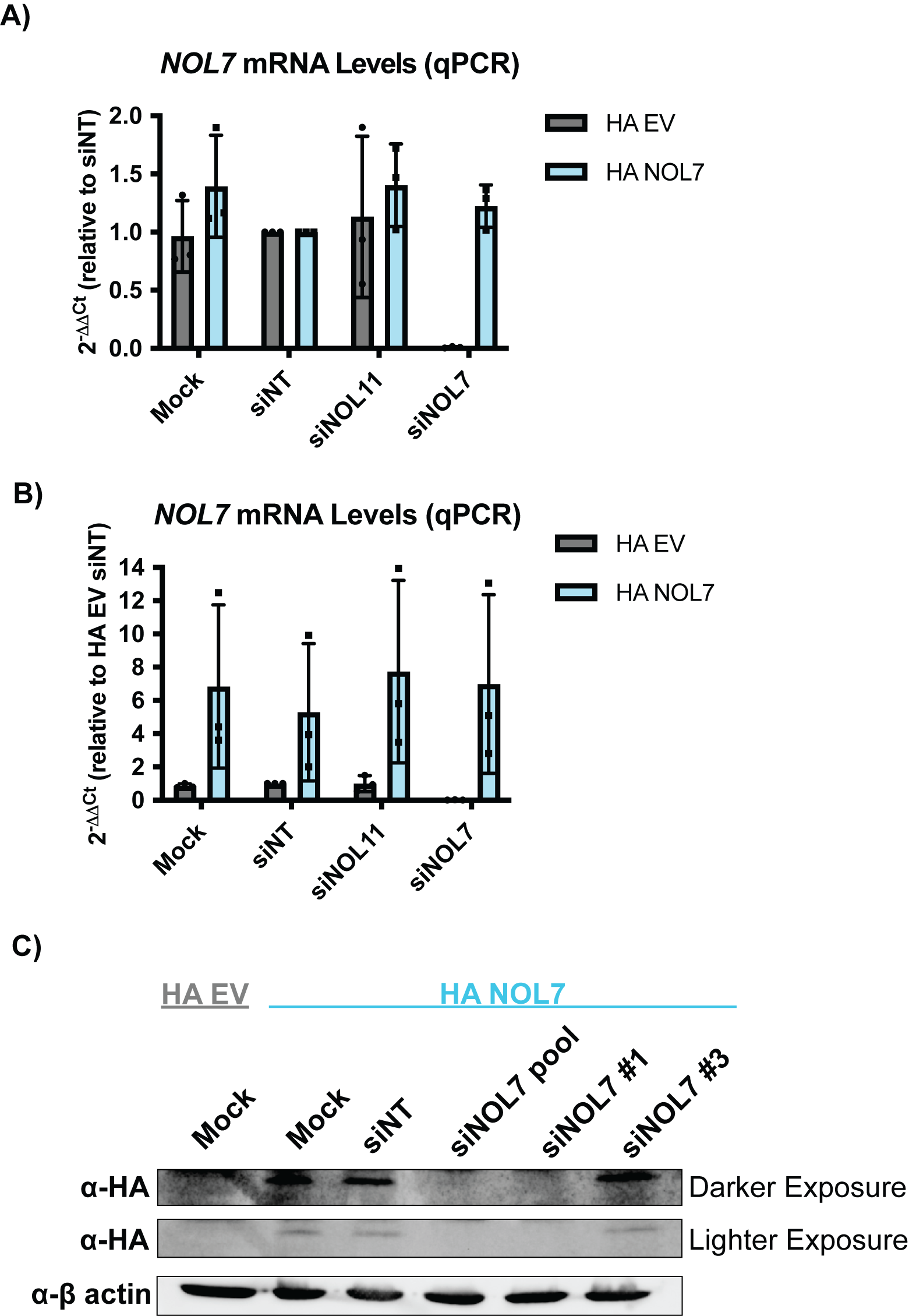

**Figure S3**: NOL7 mRNA and protein levels are rescued upon introduction of an HA-tagged and siRNA resistant version of NOL7 in HeLa cells. **A)** *NOL7* mRNA levels are rescued. qRT-PCR measuring *NOL7* mRNA levels in HeLa cells either expressing empty vector (HA EV) or siNOL7 resistant HA-tagged NOL7 (HA NOL7). 2^-ΔΔCt^ were measured relative to 7SL internal control and siNT negative control sample. siNOL7 treatment is with an individual siNOL7 #3. 3 technical replicates of 3 biological replicates, plotted mean ± SD. **B)** NOL7 mRNA levels are overexpressed in siNOL7 resistant HA tagged NOL7-transduced HeLa cells. Same as in **A)** except all samples were measured relative to siNT HA EV negative control. **C)** NOL7 protein levels are rescued only when upon individual siNOL7 #3 treatment. Western blot using an HA antibody in both HA EV and HA NOL7 transduced HeLa cells. HA EV is a negative control, while Mock and siNT HA NOL7 are positive controls. Darker exposure (top), lighter exposure (bottom), β-actin was used as a loading control.

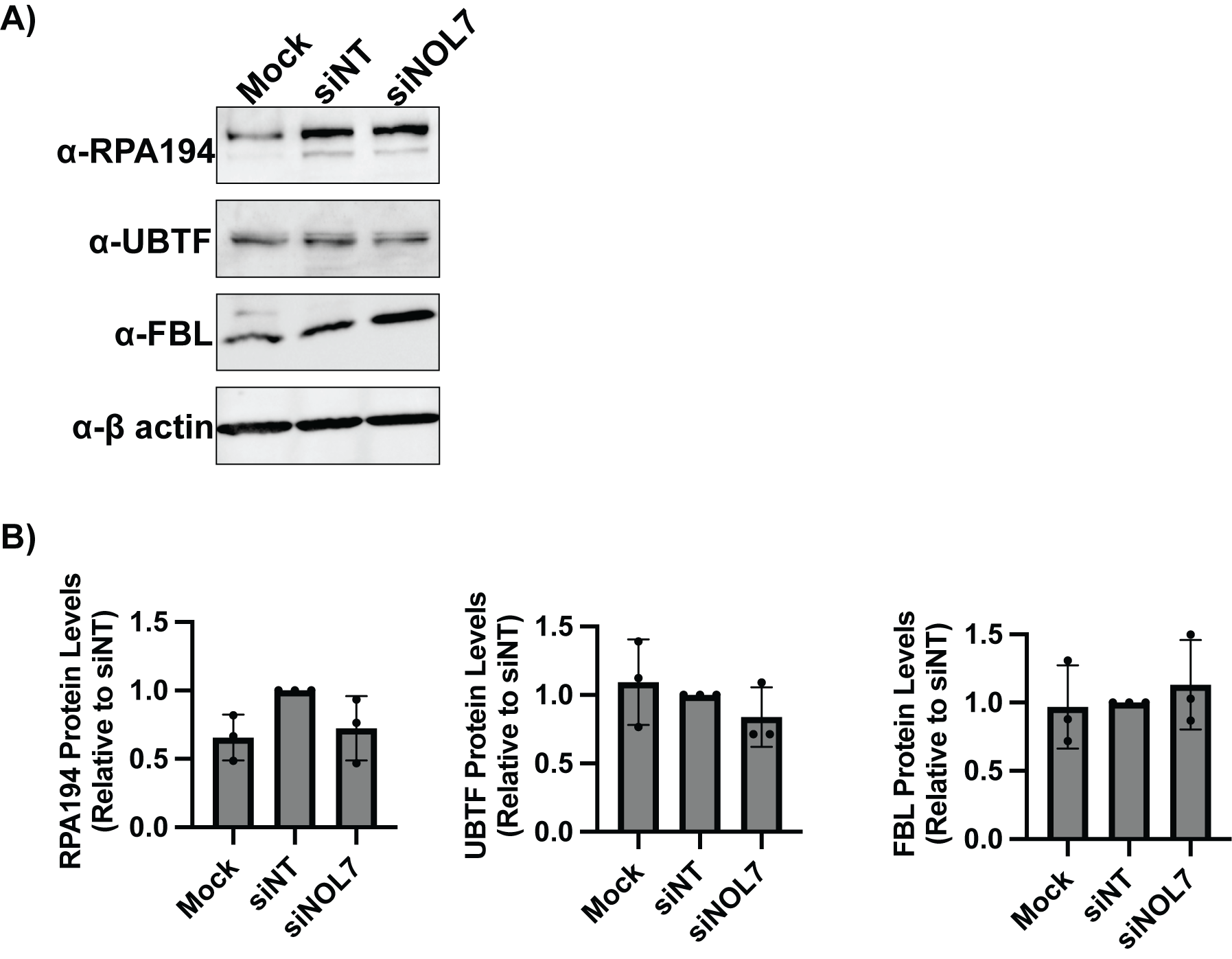

**Figure S4**: NOL7 depletion does not change the protein levels of the RNA Polymerase I transcription machinery (RPA194 and UBTF) or the methyltransferase U3 box snoRNP Fibrillarin (FBL) in MCF10A cells. **A)** Representative western blot for RPA194, UBTF, and FBL where Mock and siNT are negative controls. β-actin was used as a loading control. **B)** Quantification of western blots from **A)** normalized to β-actin signal. 3 biological replicates, plotted mean ± SD. Data were analyzed by one-way ANOVA with Dunnett’s multiple comparisons test.
